# A single mild juvenile TBI in mice leads to regional brain tissue abnormalities at 12 months of age that correlate with cognitive impairment at the middle age in male mice

**DOI:** 10.1101/2022.12.19.520991

**Authors:** Andre Obenaus, Beatriz Rodriguez-Grande, Lee Jeong Bin, Christophe J. Dubois, Marie-Line Fournier, Martine Cador, Stéphanie Caille, Jerome Badaut

## Abstract

Traumatic brain injury (TBI) has the highest incidence amongst the pediatric population and its mild severity represents the most frequent cases. Moderate and severe injuries as well as repetitive mild TBI result in lasting morbidity. However, whether a single mild TBI sustained during childhood can produce long-lasting modifications within the brain is still debated. We aimed to assess the consequences of a single juvenile mild TBI (jmTBI) at 12 months post-injury in a mouse model.

Non-invasive diffusion tensor imaging (DTI) revealed significant microstructural alterations in the hippocampus and the in the substantia innominata/nucleus basalis (SI/NB), structures known to be involved in spatial learning and memory. DTI changes paralled neuronal loss, increased astrocytic AQP4 and microglial activation in the hippocampus. In contrast, decreased astrocytic AQP4 expression and microglia activation were observed in SI/NB. Spatial learning and memory were impaired and correlated with alterations in DTI-derived derived fractional ansiotropy (FA) and axial diffusivity (AD).

This study found that a single juvenile mild TBI leads to significant region-specific DTI microstructural alterations, distant from the site of impact, that correlated with cognitive discriminative novel object testing and spatial memory impairments at 12 months after a single concussive injury. Our findings suggest that exposure to jmTBI leads to a chronic abnormality, which confirms the need for continued monitoring of symptoms and the development of long-term treatment strategies to intervene in children with concussions.

## 1. Introduction

Traumatic brain injury (TBI) has the highest incidence amongst the pediatric population [20, 63] and mild TBI (mTBI) accounts for more than 80% of all pediatric TBIs [17]. Emerging clinical data demonstrate vulnerability to long-term cognitive deficits [17]. Diffusion-weighted neuroimaging, with its potential to noninvasively detect subtle changes in tissue microstructure, is a potentially valuable tool in assessing long-term consequences of mTBI in children [74, 76] and in preclinical animal models [15, 36, 62]. These and other studies clearly demonstrate disrupted brain microstructure in vulnerable regions. However, there is still considerable debate whether a single mTBI in children can have long-lasting sequelae, whose brains are still undergoing development and reorganization at the time of injury. Clinical findings suggest that children with mTBI suffer cognitive deficits when they reach early adulthood, up to 10 years post-injury [3, 4, 11]. Despite ongoing efforts, it is still unknown whether such cognitive deficits are carried into middle to late adulthood and how the brain tissue responds cellularly and molecularly. Furthermore, brain regions may differently contribute to such long-term cognitive dysfunction after mTBI and both identification of vulnerable regions and the underlying mechanisms are not fully understood.

The importance of the hippocampus in mediating cognitive function is well established. The cholinergic system, which is diffusely localized across different brain structures, is known to play a key role in learning and memory [7]. Cholinergic projections in the basal forebrain from the substantia innominata (SI) and the nucleus basalis (NB), reach cortical and subcortical structures that regulate attention, learning and memory [2]. SI/NB atrophy was recently shown to be correlated with cognitive impairment and dementia in humans [51, 54].

To date, the relationship between the structural, molecular and behavioral long-term consequences of a single juvenile mild TBI (jmTBI) has been overlooked, especially in regard of the effects on the neurovascular unit (NVU).

To date, the relationship between the structural, molecular and behavioral long-term consequences of a single juvenile mild TBI (jmTBI) has been overlooked, especially in regard of the effects on the neurovascular unit (NVU). NVU dysfunction, such as the altered of the clearance of proteins like amyloid beta, contributes to cognitive decline involving astrocytes and the water channel aquaporin 4 (AQP4) [25, 28, 35, 50, 68]. Microglia, part of the NVU, are also known to detect vascular changes [53, 72]. Furthermore, NVU dysfunction is at the core of many progressive pathologies, including Alzheimeŕs disease [65, 67, 70]. Alterations in blood vessels [71], astrocyte phenotype [15], AQP4 expression [57], and microglial activation [31] have been reported during the first few months after juvenile mTBI (jmTBI). However, whether these cellular alterations contribute or still linger to middle age and parallel the neuroimaging changes and assocaited cognitive decline is unknown. Due to its key role in cognitive function, long-term assessment of the NVU following juvenile mTBI is critical to evaluate the long-term outcomes of chronic mTBI pathophysiology.

Based on the paucity of existing data, we explored how a single early in life mild TBI (mTBI) leads to long-term tissue-level abnormalities and associated memory dysfunction at middle age in mice. We tested our hypothesis that tissue-level diffusion MRI properties provide a non-invasive signature of ongoing gliovascular and neuronal abnormalities in the hippocampus and the basal forebrain that reflect loss of memory performance at 12 months after a single jmTBI.

## 2. Materials and Methods

### 2.1. Animals

C57BL/6J mice were bred in–house from breeder mice purchased from Janvier (Le Genest-Saint-Isle, France). TBI was administered on postnatal day (pnd) 17, only to male pups given the higher incidence of TBI in male children [16]. Mice were weaned at pnd25 and housed in groups. Animal facility was maintained at 21°C ± 1°C, 55% ± 10% humidity, in a 12h light-dark cycle and mice had access to food and water *ad libitum*. Juvenile mice were randomly assigned to experimental groups (sham and jmTBI) in weight-matched pairs. Animals that weighed below the age-appropriate range (<6g at pnd17) were excluded. Ten animals per group underwent the closed-head injury or sham procedures (jmTBI or sham), and behavioral assessment was performed at 2 and 6 months post-injury (Figure 1). At 12 months post-injury, a subset of 4 sham and 5 jmTBI mice was re-evaluated for behavior and ex vivo and immunohitochemestry experiments were performed (Figure 1). All animal procedures were carried out in accordance with the European Council directives (86/609/EEC) and the ARRIVE guidelines.

**Figure 1:**
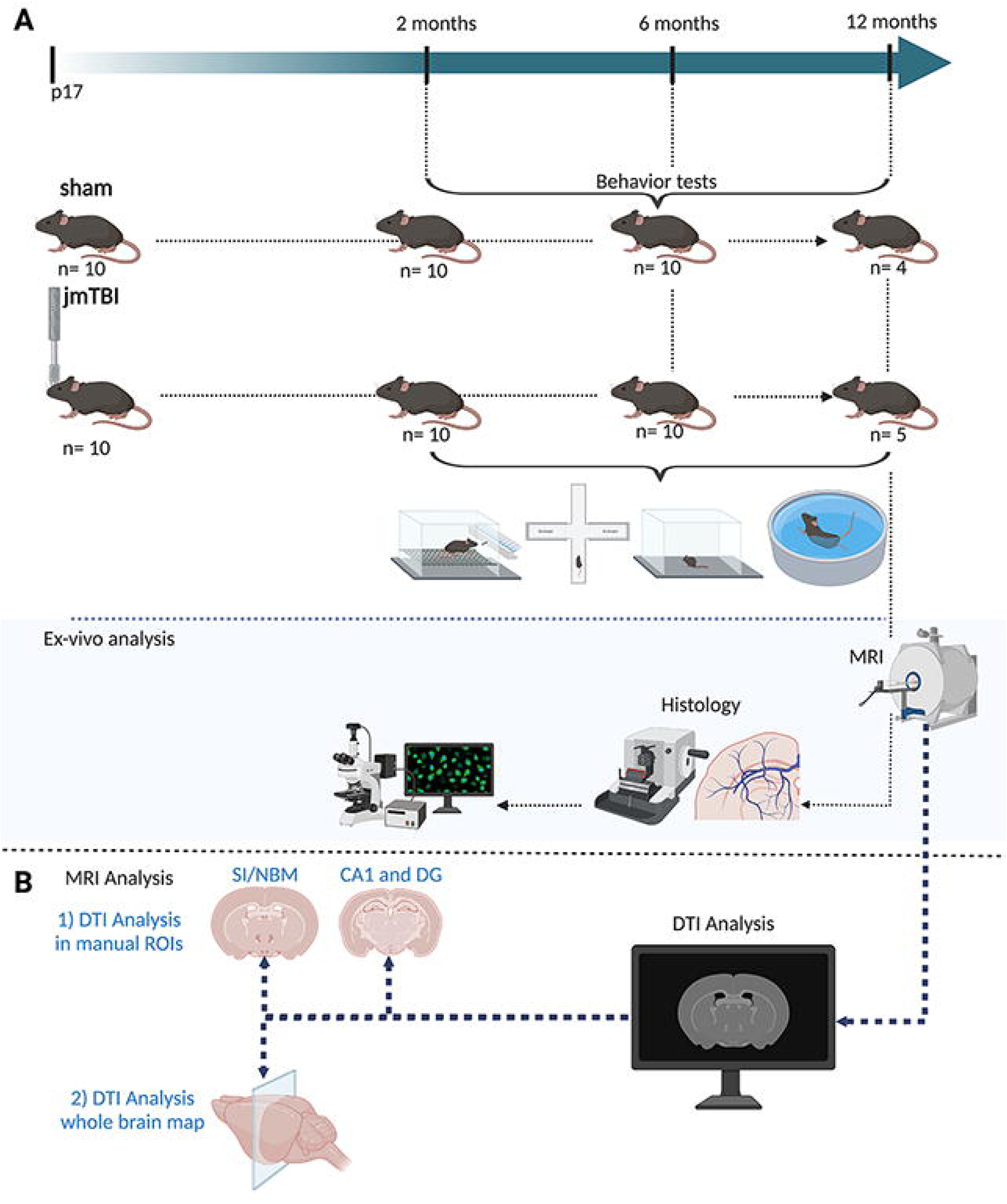
Experimental design. A. Schematic illustration of the temporal experimental design with repeated behavior evaluations in sham and jmTBI mice followed by high resolution diffusion *ex vivo* MRI and histological analyses. B. MRI Analysis has been carried out with 1) manual ROIs and 2) Whole brain map

### 2.2. Closed-Head Injury with Long-term Disorders (CHILD)

Closed head injury model with focal injury, head acceleration, and rotational component was performed at pnd17 as previously described [57]. Briefly, mice were anaesthetized using 2.5% isoflurane and 1.5 l/min air for 5 min and impacted directly over their intact head using an electromagnetic impactor (Leica Impact One Stereotaxic impactor, Leica Biosystems, Richmond, IL, USA) with a 3mm round tip at a speed of 3m/s, a depth of 3mm and a dwell time of 100ms. The tip of the impactor was placed over the left somatosensory-parietal cortex center at ∼Bregma 1.7 mm. Sham mice underwent the same anesthesia procedure and were placed under the impactor, but did not receive any impact. Mice were allowed to fully recover in an individual cage before being reintroduced to their home cage.

### 2.3. Behavioral Assays

Behavioral tests were performed at 2, 6 and 12m post-TBI (13m of age). Tests were video-recorded (Fujifilm Camera, Imaging Source, Germany) and tracking was performed using ANY-maze (Stoelting Co., Italy).

#### 2.3.1. Open field test

Mice were initially positionned in the center of the open field (45 × 45 cm, 35 cm height) and allowed to freely explore the environment for 5 min. Between sessions, the open field was cleaned with a 50% EtOH solution. Mean speed, total distance travelled and time spent in the center of the arena (defined as the 35 × 35 cm central part) were measured to estimate general motor activity and anxiety-like behavior.

#### 2.3.2. Elevated plus maze test

Mice were positionned on the center square of a gray polypropylene elevated plus maze (66 x 66 cm, 6 cm wide arms, walls of 15 cm height, elevation of 40 cm above the floor) facing an open arm and allowed to freely explore the apparatus for 5 min. Time spent in the open and closed arms were scored.

#### 2.3.3. Sucrose preference test

Mice were single-housed in a typical home-cage (50 × 25 × 20cm) and their consumption of plain or sweet (5% sucrose, Sigma-Aldrich, Saint-Quentin-Fallavier, France) water was quantified over 24 hours. The preference ratio (PR) was defined as the sucrose intake divided by the total (water + sucrose) intake.

#### 2.3.4. Morris water maze (MWM) test

The test was performed as detailed previously [30]. Briefly, a tank of 100 cm diameter filled with opaque water and a 10 cm diameter platform was used. On the first day a cued test was performed, in which the platform was 2 cm over the level of the water and had a protruding rod which made the platform visible. Animals were placed in the tank and required to swim to the visible platform to end the task. Each animal completed 5 blocks of 2 trials. Platform position changed in each block, the starting location was shifted in each trial.

On day 2, a spatial learning test was performed (spatial learning, SL). Each animal completed 5 blocks of 2 trials in which the starting position changed through the test. The platform remained in a fixed position during the whole test, submerged 2 cm under the opaque water. After 24h (day3), a memory probe test was performed (P). The platform was removed and mice were released from the opposite side of the water maze and left to swim for 60 sec. For the cued and learning tests, if mice had not found the platform after 60 sec, they were guided to it and left to remain on it for 5 seconds. Cumulative distance (sum over time of the mouse’s distance to the platform as measured 5 times every second) to the platform was used as the outcome variable during the cued and learning tests. The percentage of time in the platform quadrant was used as the outcome variable during the probe tests.

#### 2.3.5. Novel location test

Mice were first habituated for 5min to the empty open field described above with a green tape landmark on one of the walls . Then mice were presented with two identical objects placed on a side of the open field for 10 min, as previously described [42]. Four hours later, one of the objects was moved towards a diagonal position and the mouse was reintroduced and left to explore for another 10 minutes. Between session, the open field and objects were cleaned with a 50% EtOH solution. A discrimination index (DI) was calculated as the difference of the time spent exploring the object in the new location minus the old location over the total time exploring both objects.

### 2.4. Magnetic Resonance Imaging (MRI) and Analysis

After the final behavioral tests, mice were anesthetized (ketamine/xylazine, 100mg/kg and 20mg/kg respectively) and perfused transcardiacly with a 4% paraformaldehyde solution (PFA, in phosphate-buffered saline, PBS). Brains were kept in 4% PFA overnight at 4°C, and then transferred to PBS with 0.1% sodium azide and stored at 4°C. For *ex vivo* MRI, brains were immersed in fluorinert (Sigma-Aldrich, Saint-Quentin Fallavier, France) and scanned using a 7.0T scanner (Bruker BioSpin, Billerica, MA). Supplementary Table 1 contains the detailed sequence parameters that were utilized. *Ex vivo* imaging was used as it allows for increased resolution, improved signal to noise ratios, and reduced motion artifacts.

Parametric maps for T2 were processed using the MRI Processor plugin from FIJI (NIH, USA) [59, 60]. All DTI quality control, image processing, and region of interest (ROI) delineations were performed using DSI studio (http://dsi-studio.labsolver.org, Ver: 2.0 Department of Neurological Surgery, University of Pittsburgh, USA). All acquisitions were examined for motion and ghosting artifacts were examined in each direction of the acquisition and mice were discarded if visible motion was observed in greater than four directions. Eddy current corrections were applied and DTI parametric maps of fractional anisotropy (FA), axial diffusivity (AD), mean diffusivity (MD) and radial diffusivity (RD) were obtained. Manual regional delineation of the DG, CA1 and SI/NB was performed at two Bregma levels, -1.5mm for the hippocampus and Bregma 1.5mm for the substantia innominate/ Nucleus Basalis (SI/NB). All data analysis was undertaken by a blinded experimenter.

Automated whole-brain regional analysis of hippocampus and basal forebrain components were performed as follows: T2 images were N4 bias field corrected with Advanced Normalization Tools (ANTs v2.1) and extra cranial tissues were masked using 3D Pulse-Coupled Neural Networks (PCNN3D v1.2) [5, 14, 55, 69, 73]. The mean of B0s from the DTI acquisition underwent the identical procedures. T2 data were then registered using ANTs Symmetric Normalization (SyN) algorithm to match the DTI, after which the same algorithm was utilized to register a custom-made atlas and labels based on Australian Mouse Brain Mapping Consortium (AMBMC) to the same native space [55, 69]. The transformation was applied to the label maps for 3-dimensional segmentation. The resulting label maps encompassed the entire brain, so that region segmentations would cover the entire length of a given structure over multiple image slices. The T2 relaxation times (ms) and volumetric data were extracted based on the final curated segmentation. All diffusion data also underwent eddy correction and was reconstructed using FMRIB Software Library’s DTIFIT where DTI metrics were extracted using the transformed label maps (FSL v5.0; FMRIB, Oxford, UK) [5, 73]. FA, MD, RD and AD diffusivity maps were used for final data acquisition. Hippocampal subregions (CA1, CA2, and CA3) and basal forebrain cholinergic circuit components (SI, substantia innominate (including ventral palladium); NB, nucleus basalis; MS, medial septal nucleus) were examined in detail.

### 2.6. Immunohistochemistry and Analysis

Brains were rinsed in PBS after MRI scanning and were cut into 50μm-thick coronal sections using a vibratome (Leica, Richmond, IL). Sections were stored at -20**°**C in cryoprotective solution (30% ethylene glycol and 20% glycerol in PBS) until further use. For immunohistochemical staining, brains were washed in PBS. Basic antigen retrievals were performed for Iba1-immunolabeling with a 30 min incubation in 10mM Sodium Citrate at 80**°**C after which sections were allowed to reach room temperature (RT) in the same solution. After antigen retrieval, sections were extensively washed in PBS. Sections were then incubated in blocking solution (1% BSA, 0.3% Triton X-100 in PBS) for 1h at RT followed by overnight incubation at 4**°**C with primary antibodies diluted in blocking solution. Sections were washed in PBS and then incubated for 2h at RT with the corresponding fluorescent secondary antibodies diluted in blocking solution. Primary antibodies were chicken polyclonal anti-mouse GFAP (1:500, Millipore, Billerica, MA), rabbit polyclonal anti-mouse AQP4 (1:300, Chemicon International, Temecula, CA, USA), Iba-1 (1:200, Abcam, Cambridge, MA), NF200 (1:500, Chemicon International, Temecula, CA, USA) rabbit monoclonal amyloid precursor protein (APP) antibody [Y188] (1:500, Abcam, Cambridge, MA) and NeuN (1:500, Abcam, Cambridge, MA). All secondary antibodies had Alexa-Fluor dyes (Molecular Probes, Invitrogen, Carlsbad, CA) and were used diluted 1:1000. Fluorescein-conjugated tomato lectin (1:500, Vector, Vector Laboratories, Burlingame, CA) was used to detect vessels.

Brains were mounted onto glass slides and cover-slipped using Vectashield (Vector, Vector laboratories, Burlingame, CA) with DAPI (1/10000, thermoFisher scientific). Slides were kept at 4**°**C. Slides were scanned at 10x magnification using a slide-scanner (Hamamatsu Nanozoomer 2.0HT, Bordeaux Imaging Center, Bordeaux, Fr). Confocal images were taken at 40x magnification using a Nikon Eclipse Ti inverted laser scanning microscope with a Nikon acquisition software (NIS-Element, NIS). Settings were kept constant amongst experimental groups within each batch of experiments. Antibodies are diffusing through the section approximatively on 10 μm thickness on each side of section. Hence, the confocal acquisitions were done on 10μm thickness corresponding to the part of the tissue closer to the surface.

Image analysis was performed using FIJI software (Schindelin, 2012, Fiji: an open-source platform for biological-image analysis) in a blinded manner on 2 brain sections containing the region of interest hippocampus and SI/NB in the ispi-and contralateral hemisphere to the impact. For each region, 3 different images were taken either at 20X or 40X using a confocal microscope (Nikon Eclipse Ti inverted laser scanning microscope). Given the irregular anatomy of the SI/NB, the region was manually delineated within each section to obtain values within the region of interest.

For quantification of the APP staining in the hippocampus, APP images were acquired at 20X using Nikon Eclipse Ti inverted laser scanning microscope with a Nikon acquisition software (NIS-Element, NIS). Using FIJI software, stack images were reconstructed using Z-stack function. The resultant stack was first treated with “Bandpass” filter without autoscale after filtering, followed by “Unsharp mask (Radius: 1.0 pixel – Mask Weight: 0.9). Filtered images were then binarised after thresholding (6-7% to delineate the thin processes). Then the images were skeletonized and particle analysis in the CA1 and stratum oriens was perform. The size the particles were expressed as surface area staining. The process analysis is summarized in supplementary figure 1.

For quantification of AQP4 in perivascular endfeet and astrocytic processes, GFAP-AQP4 double staining was performed and confocal images with 40X lens (Nikon) were acquired. Using FIJI software, astrocytes were manually outlined in each image in the GFAP channel, and that region of interest was then copied into the AQP4 channel, from which the mean OD value was obtained. For the measurement of perivascular staining, vessel-like (tubular) structures were manually outlined and mean OD was measured in this region.

For the quantification of the percentage of the tomato lectin (TL)-positive areas, images were manually thresholded using the FIJI threshold tool to enhance the vessels from background on the 3 images taken at 40X from each brain region. For the quantification of vascular diameter in TL-stained sections, the diameter was measured manually using the straight-line FIJI tool from 8-10 vessels per image with a total of n= 177 ± 23 blood vessels.

For the quantification of morphological features of microglia, skeleton analysis was performed on 3 different pictures were taken with 20X in 2 adjacent brain sections for each ROIs (protocol adapted from [15, 38]). Briefly, automated background correction and manual thresholding were applied to image stacks to remove background before binarizing and skeletonizing the images using FIJI. Small non-cellular structures corresponding to isolated processes or background spots were filtered out by the following criteria: number of branches > 4, number of endpoints> 4, number of junctions > 3. Number of branches, number of slab voxels that correspond to branch length and number of endpoints were kept as parameters of interest provided by skeleton analysis. The analyses were carried out on 45-58 microglia from n=4 shams and n=5 jmTBI mice from 2 brain section levels.

### 2.7. Statistical analysis

GraphPad Prism 3.05 (GraphPad Software Inc., USA) was used for statistical analysis. For comparison of single features between two groups, an unpaired t-test was used. For comparisons of several features or a feature across several regions of interest, a two-way ANOVA followed by Tukey post-hoc test was used. Linear regression was applied to assess correlations between two parameters. Significance was reported at p < 0.05. All data are presented as the mean ± standard error of the mean (SEM) and followed a normal distribution (Shapiro-Wilk test). Whole-brain analysis of T2 and DTI data were analyzed with GraphPad Prism 7.00 using an unpaired t-test with Welch’s correction for each ROIs. All MRI data were examined for sample outliers using 1.5 interquartile range (IQR) testing.

## Results

### Brain tissue alterations on MRI

Examination of the T2 images showed no structural changes at 12m after a single jmTBI nor alterations in T2 relaxation values (Figure 2A). Manual ROI delineations found significant decreases in FA within the ipsilateral hippocampal dentate gyrus (DG) of jmTBI mice compared to shams was observed (figure 2B, C). In the ipsilateral but not contralateral SI/NB, AD was also significantly decreased (figure 2D). Other DTI scalars (MD, RD) were not found to be significantly altered within hippocampal and SI/NB at 12 months after jmTBI (see Supplementary Table 2 for data and inter-group comparisons).

**Figure 2:**
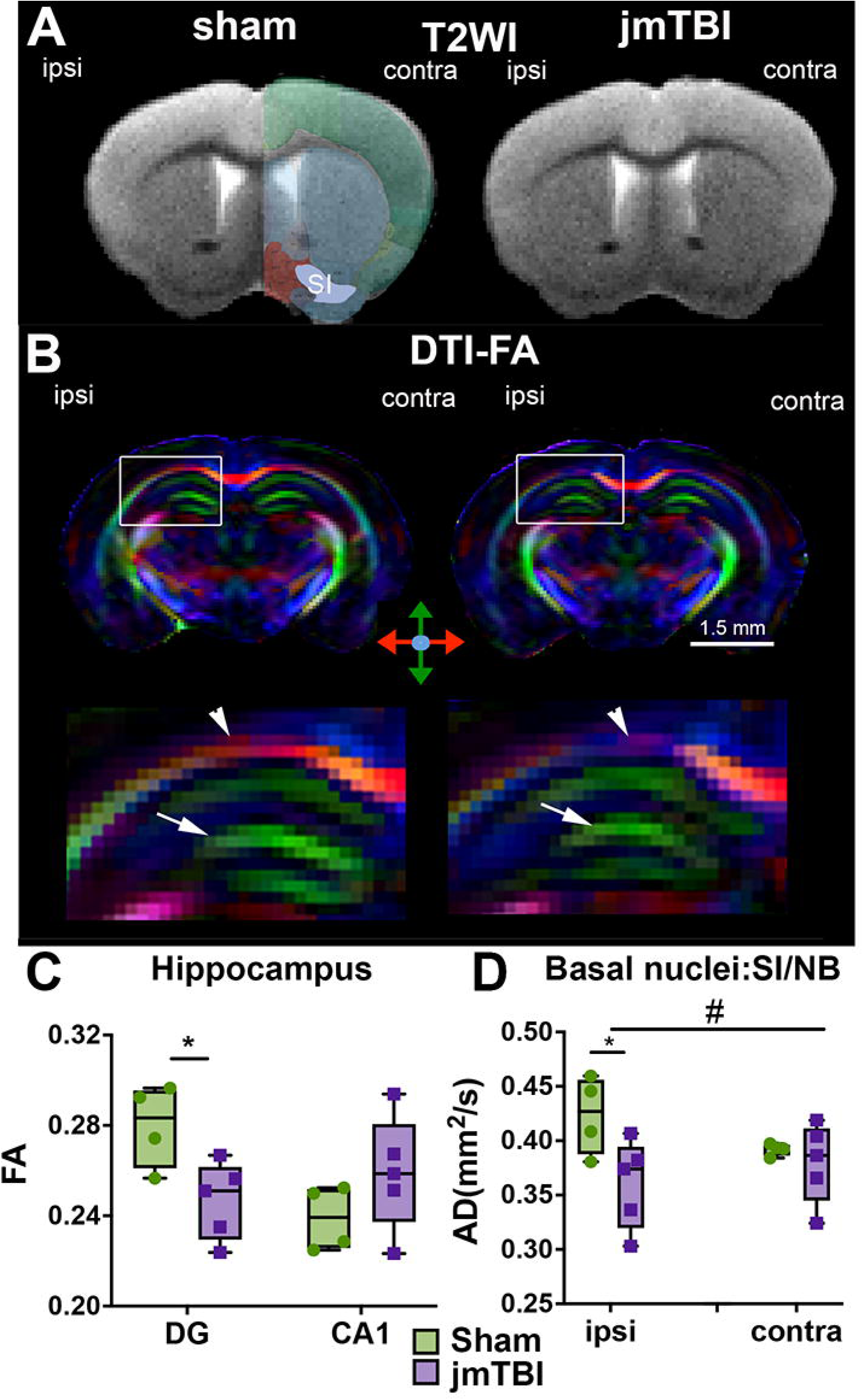
DTI alterations are detected in the hippocampus and SI/NB 12m after jmTBI. A. Representative T2WI images of sham and jmTBI mouse brains at the coronal level of the SI/NB indicated on contralateral hemisphere. T2WI did not show gross morphology difference between sham and jmTBI. B. Representative coronal pseudo-colored fractional ansiotropic (FA) images of sham and jmTBI mouse brains at the site of the impact (top panels) with expanded FA maps of the ipsilateral hippocampus (bottom panels). Arrowhead highlights altered FA within the corpus callosum and arrohead indicate the CA1 region of the hippocampus. C. FA was significantly decreased within the hippocampal DG but not CA1. D. AD was significantly decreased of in the ipsilateral but not contralateral SI/NB. C. and D. Two-way ANOVA (# indicating global jmTBI vs sham differences) with Sidack post-hoc test (* indicating jmTBI vs sham). Data expressed as mean+SEM. # or *P<0.05

To further examine microstructural alterations within the hippocampus and basal forebrain, we utilized whole-brain segmentation methods based on our modified AMBMC atlas to analyze hippocampal subregions (CA1, CA2, and CA3) and basal forebrain cholinergic circuit components (SI, substantia innominate; NB, nucleus basalis; MS, medial septal nucleus) over the entirety of the given structure in 3D (Table 1). Using this atlas-based approach, we found that the contralateral CA2 exhibited a significant increase in T2 values (*p=0.0442, t=2.926, df=3.912) but other hippocampal regions showed no significant differences in any of the imaging metrics. Trending increases were noted in the ipsilateral CA2 in MD and RD (MD: #p=0.069, t=2.255, df=5.493; RD: #p=0.0626, t=2.295, df =5.855). Basal forebrain regions exhibited significant differences between 12 months old sham and jmTBI mice where the ipsilateral SI had significant decreases in FA (*p=0.0068, t=5.044, df=4.11). Both the ipsilateral and contralateral NB regions exhibited tissue level decreases in RD (ipsilateral: *p=0.0394, t=2.536, df=6.892; contralateral: *p=0.0384, t=2.933, df=4.356) while only contralateral NB tissue exhibited MD differences (*p=0.0365, t=2.596, df=6.80, Table 1). Thus, 12 months after a single jmTBI there are brain regions that exhibit significant MRI-derived microstructural alterations.

**Table 1.** T2WI and DTI reveal tissue-level alterations 12m following CHILD (t-values)

| Region | Ipsilateral |  |  |  |  | Contralateral |  |  |  |  |
| --- | --- | --- | --- | --- | --- | --- | --- | --- | --- | --- |
|  | T2 | FA | MD | AD | RD | T2 | FA | MD | AD | RD |
| Hippocampus |  |  |  |  |  |  |  |  |  |  |
| CA1 | 1.10 | 0.19 | 0.37 | 1.15 | 0.37 | 0.49 | 0.94 | 0.31 | 0.06 | 0.99 |
| CA2 | 1.62 | 1.42 | **2.26<sup>#</sup>** | 2.05 | **2.30<sup>#</sup>** | **2.93<sup>\*</sup>** | 1.72 | 0.27 | 0.47 | 0.18 |
| CA3 | 1.64 | 0.44 | 1.36 | 1.01 | 1.24 | 0.10 | 0.35 | 0.14 | 0.36 | 0.02 |
| DG | 0.85 | 1.99 | 0.86 | 0.26 | 1.07 | 0.19 | 0.78 | 1.33 | 0.29 | 1.53 |
| Basal Forebrain |  |  |  |  |  |  |  |  |  |  |
| SI | 0.12 | **5.04<sup>\*</sup>** | 1.92 | 1.69 | 1.34 | 0.70 | 0.34 | 0.49 | 0.76 | 0.23 |
| NB | 0.81 | 1.89 | 0.41 | 0.14 | **2.93<sup>\*</sup>** | 0.75 | 1.52 | **2.60<sup>\*</sup>** | **2.43<sup>#</sup>** | **2.54<sup>\*</sup>** |
| MS | 0.72 | 0.6 | 0.76 | 0.17 | 0.38 | 0.40 | 0.63 | 0.11 | 0.10 | 0.10 |

Note. Sham vs. CHILD. T2 (ms), MD (mm<sup>2</sup>/s), AD (mm<sup>2</sup>/s), RD (mm<sup>2</sup>/s).
DG, dentate gyrus; SI, substantia innominate (ventral palladium); NB, nucleus basalis; MS, medial septal nucleus. <sup>\*</sup>*p*<0.05, <sup>#</sup>*p*<0.1

### Neuronal alterations in the hippocampus and SI/NB

To identify neuronal changes, we utilized NeuN immunohistochemistry to identify neuronal cell bodies, NF200 immunostaining to reveal axons and dendrites and APP immunolabeling present in neurons. A significant decrease in NeuN labeling was found within the ipsilateral DG and CA1 (Figure 3A,B) but not in the contralateral regions (Figure 3B). Similarly, APP staining showed neuronal process fragmentation in the stratum radiatum after jmTBI (Figure 3C) which was confirmed by a significant decrease in surface area staining in the injured group (Fig. 3D). NF200 staining was not significantly different between experimental groups in the hippocampus (data not shown), but was significantly increased in the contralateral SI/NB in jmTBI compared to sham mice (figure 3E, F). However, there was a correlation between decrease of NF200 and FA values in CA1 (see supplementary Figure 3). Unlike the hippocampus, there were no group differences in NeuN staining in the SI/NB (data not shown).

**Figure 3:**
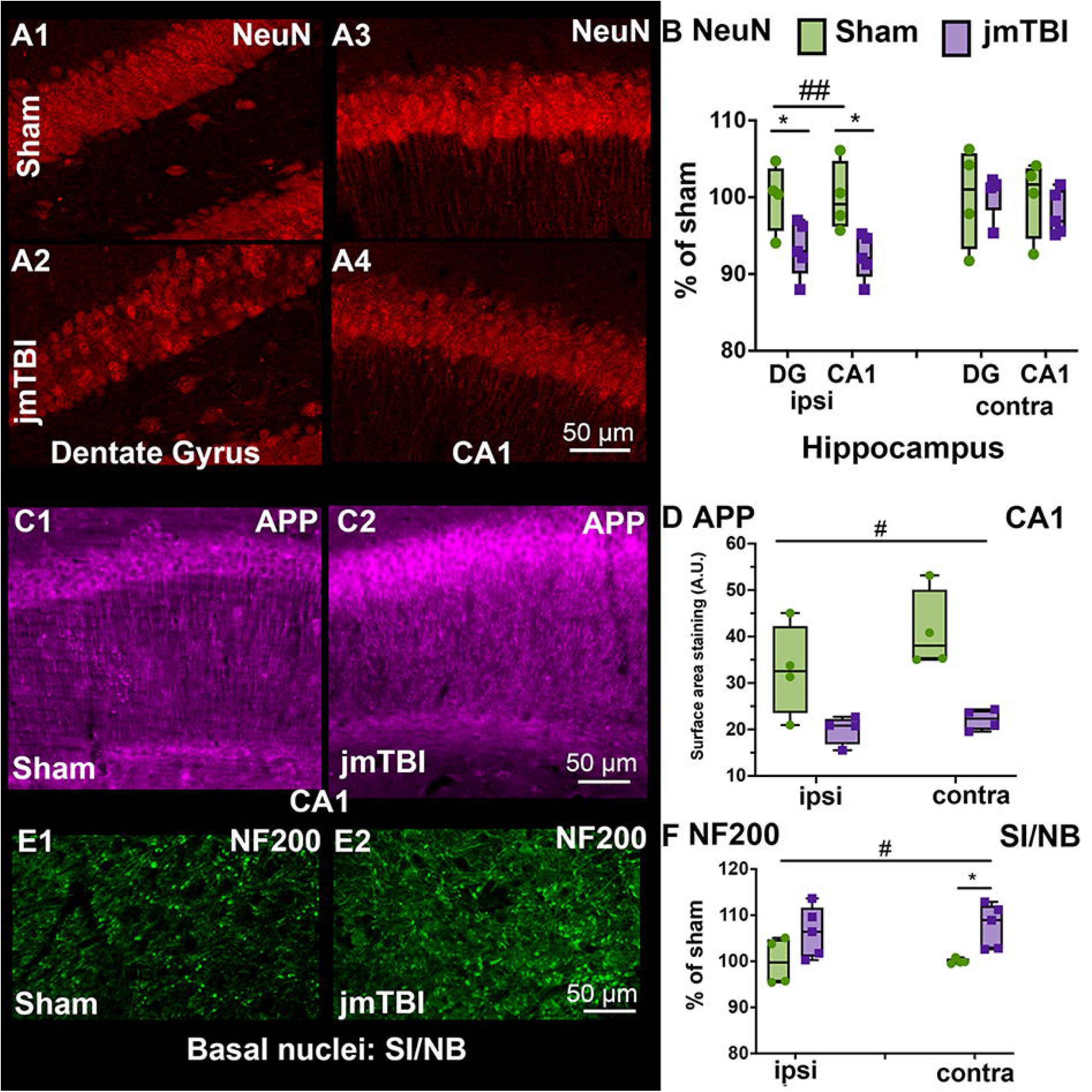
jmTBI leads to hippocampal neurodegeneration and SI/NB axonal alterations at 12mo post injury. A. Decreased NeuN (neuronal) density as a marker of neurodegeneration was observed in the DG (A2) and ipsilateral CA1 (A4) of jmTBI compared to sham mice (A1 and A3). B. A significant decrease in NeuN staining in the ipsilateral but not the contralateral regions of the hippocampus of jmTBI mice was observed. C. APP staining in the ipsilateral CA1 region of sham (C1) and jmTBI (C2) mice revealed a fragmented staining patterns in stratum radiatum under the CA1 pyramidal layer in jmTBI (C2) compared to sham group (C1). Increased staining was observed in the CA1 pyramidal layer of jmTBI (C2) mice. D. APP staining quantification showed an overall decrease in surface area staining in the jmTBI group. E. Neurofilament NF200 staining was observed in the SI/NB of sham (E1) and jmTBI (E2). The intensity of the staining is higher in jmTBI group than sham mice (E1) F. NF200 immoreactivity was significantly increased between groups (#) with significant differences found between sham and jmTBI on the contralateral side (*). B, D, F, Two-way ANOVA (# indicating global jmTBI vs sham differences) with Sidak post-hoc test (* indicates jmTBI vs sham differences in the DG or CA1 in B and C; and in the ipsilateral or contralateral in D and F). Data expressed as mean+SEM. # or *P<0.05, ## P<0.01. (passed Shapiro-Wilk normality test)

### Differential glial transformations in the hippocampus and SI/NB

To characterize the changes within the glial compartment, GFAP and AQP4 labeling were used to evaluate astrocytic changes, and IBA1 immunolabeling was performed to assess microglia. In addition, tomato lectin (TL) staining was used to identify blood vessels and their relationship to astrocytes. Reductions in GFAP immunostaining were observed in the contralateral SI/NB (figure 4 A, B). It is paralleled by an increase of the number of blood vessels with small diameters under 5μm measured from TL staining in jmTBI animals compared to sham (figure 5 and supplementary figure 2). In contrast, GFAP immunostaining remained unaltered in the ipsilateral SI/NB and in ipsi and contralateral sides of the hippocampus (figure 4A).

**Figure 4:**
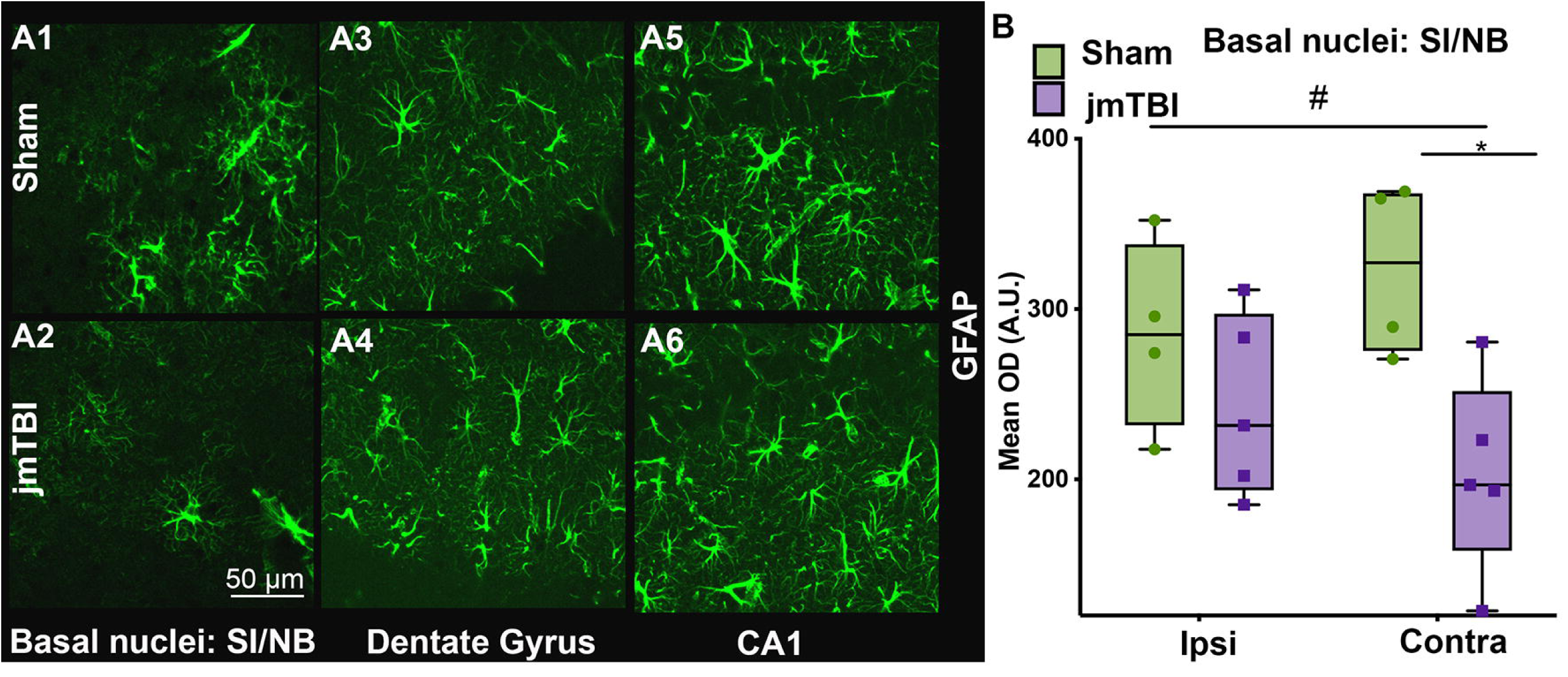
Astrogliosis is reduced in the SI/NB 12m after jmTBI but not within the hippocampus. A. A reduction in GFAP staining was observed in the SI/NB of jmTBI mice (A2) compared to sham mice (A1) but no overt observed differences in the hippocampus (A3-A6). B. GFAP decreases were significantly decreased on the contralateral but not ipsilateral side. Two-way ANOVA (# indicating global jmTBI vs sham difference) with Sidak post-hoc test. Data expressed as mean+SEM. # and * P<0.05 (passed Shapiro-Wilk normality test)

**Figure 5:**
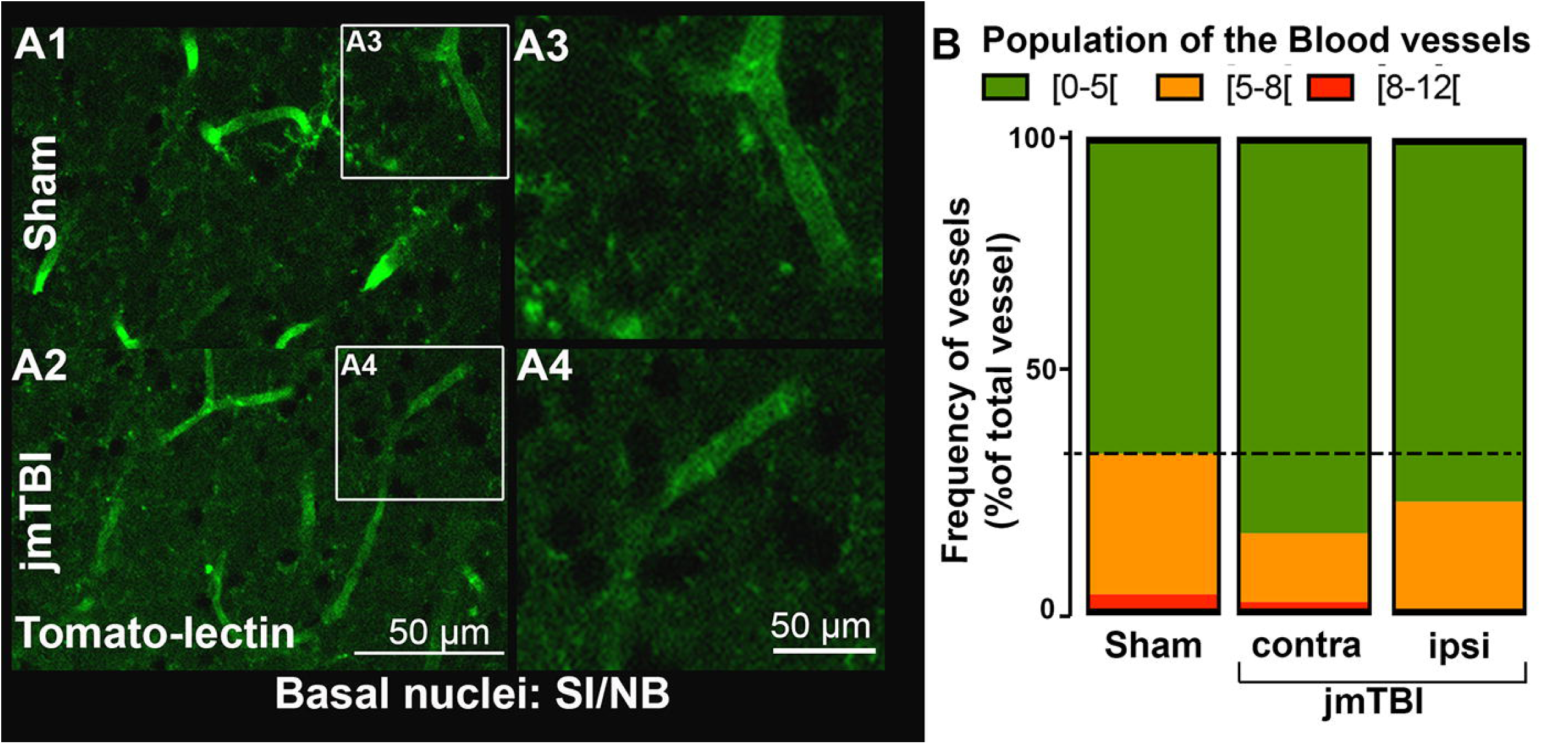
Vascular change was observed in the SI/NB 12m after jmTBI. A. Blood vessels were stained with tomato-lectin in both groups. A decrease in vessel diameter was observed in the SI/NB of the jmTBI (A2, A4) compared to sham group (A1, A3). B. Significant increase of the proportion of small blood vessel with diameter between [0-5μm[in jmTBI compared to sham mice in both ipsilateral and contralateral sides of the SI/NB.

AQP4 was expressed in astrocytic perivascular endfeet and processes with notable differences between groups and brain structures (figure 6). Subcellular quantification of AQP4 staining in perivascular structures (adjacent to blood vessels) and astrocyte processes displayed decreased AQP4 expression levels in both ipsilateral and contralateral SI/NB (figure 6A, B). In contrast, AQP4-staining in both ipsilateral and contralateral DG was increased in astrocytic perivascular and processes in jmTBI compared to sham mice (figure 6 C,D). In the ipsilateral CA1 (but not contralateral) perivascular AQP4 was also increased, with no overt change in AQP4 in astrocytic processes (figure 6 E,F). Hence, jmTBI results in differential astrocytic AQP4 expression in a region-specific manner at 12 months after injury.

**Figure 6:**
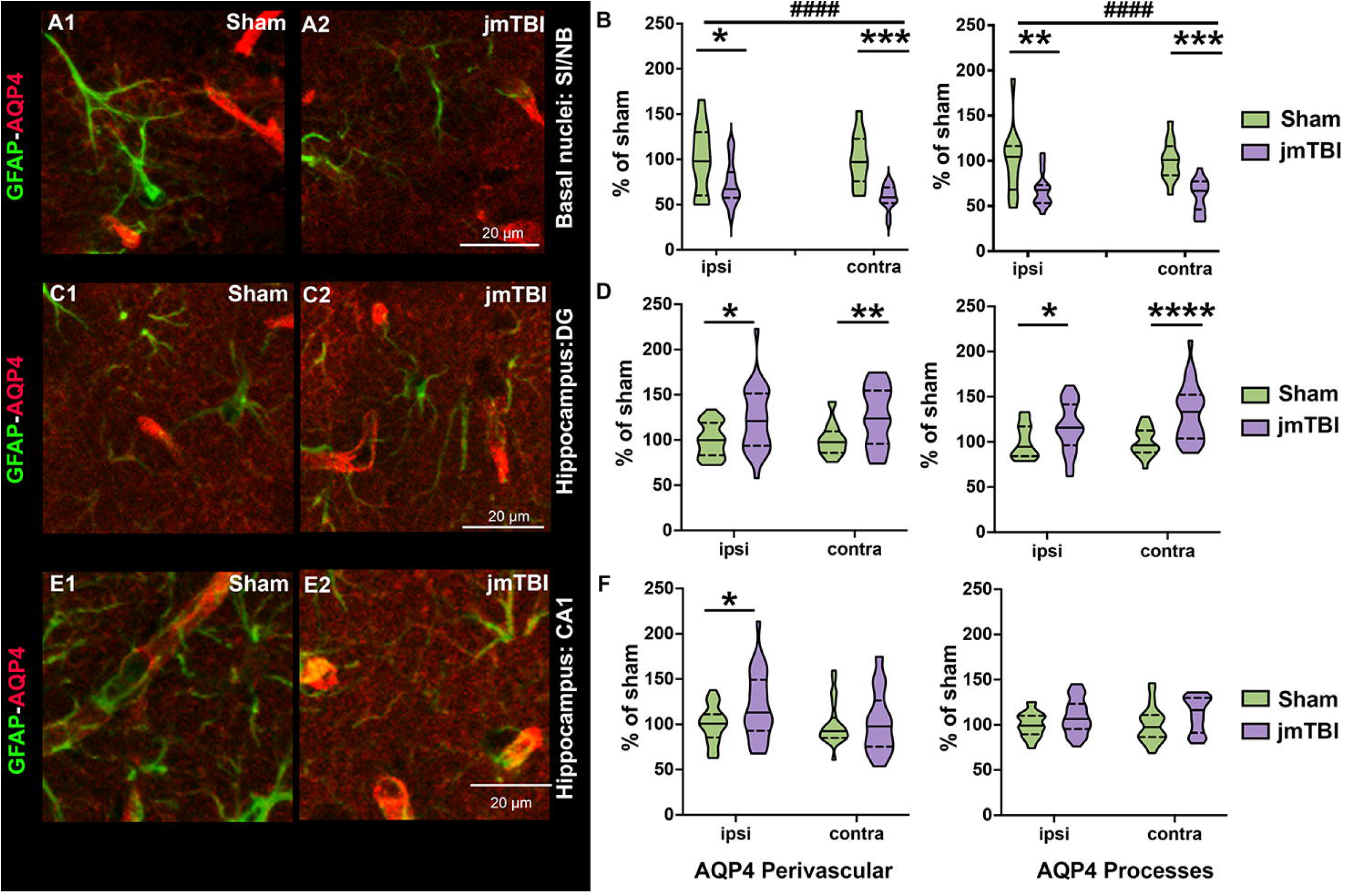
AQP4 relocalizes to alternate cellular locations in the SI/NB and hippocampus 12m after jmTBI. A. GFAP-AQP4 double-staining in representative images of perivascular structures corresponding to astrocyte endfeet in contact with blood vessels and astrocyte processes in the SI/NB . B. AQP4 quantification within perivascular astrocyte endfeet (left graph) and astrocyte processes (right graph) revealed significant AQP4 reductions in the SI/NB. Two-way ANOVA (# indicates global jmTBI vs sham differences) with Sidack post-hoc test (* indicates jmTBI vs sham differences in the ipsilateral or contralateral side). Data expressed as mean+SEM. *P<0.05, **P<0.01, ***P<0.001, ####P<0.0001 C. Representative images of perivascular structures corresponding to astrocyte endfeet in contact with blood vessels and astrocyte processes in the hippocampal DG from GFAP-AQP4 double stained sections. D. AQP4 staining was significantly increased in DG perivascular astrocyte endfeet (left graph) and astrocyte processes (right graph) in the ipsilateral and contralateral regions. E. GFAP-AQP4 staining in the hippocampal CA1 perivascular structures corresponding to astrocyte endfeet in contact with blood vessels and astrocyte processes. E. AQP4 staining was significantly increased in perivascular structures (left graph) in the ipsilateral CA1 of jmTBI mice. There were no differences in AQP4 from contralateral perivascular structures alongside no changes within astrocytes. Two-way ANOVA (# indicates global jmTBI vs sham difference) with Sidack post-hoc test (* indicates jmTBI vs sham differences in DG or CA1 subregions of the hippocampus). Data expressed as mean+SEM. # or *P<0.05, ## or **P<0.01, ###P<0.001, #### or ****P<0.0001 (passed Shapiro-Wilk normality test)

Microglial (IBA1) staining was significantly increased in the contralateral SI/NB in jmTBI compared to sham mice (Figure 7A, B). No change in IBA1 staining density was found in hippocampal DG and CA1 (data not shown). The increased IBA1 staining in the contralateral SI/NB was reflected by a decrease in branch lengths measured using morphological-skeleton analysis (Figure 7C, D). While IBA1 staining density was not altered in the CA1 we found decrements in branch length, number of branches and number of endpoints (Figure 7E, F), consistent with morphological alterations. Microglia in jmTBI mice exhibited a more retracted shape than in shams, suggesting that activated microglia are still present 12 months post-injury, mostly contralateral to the injury site.

**Figure 7:**
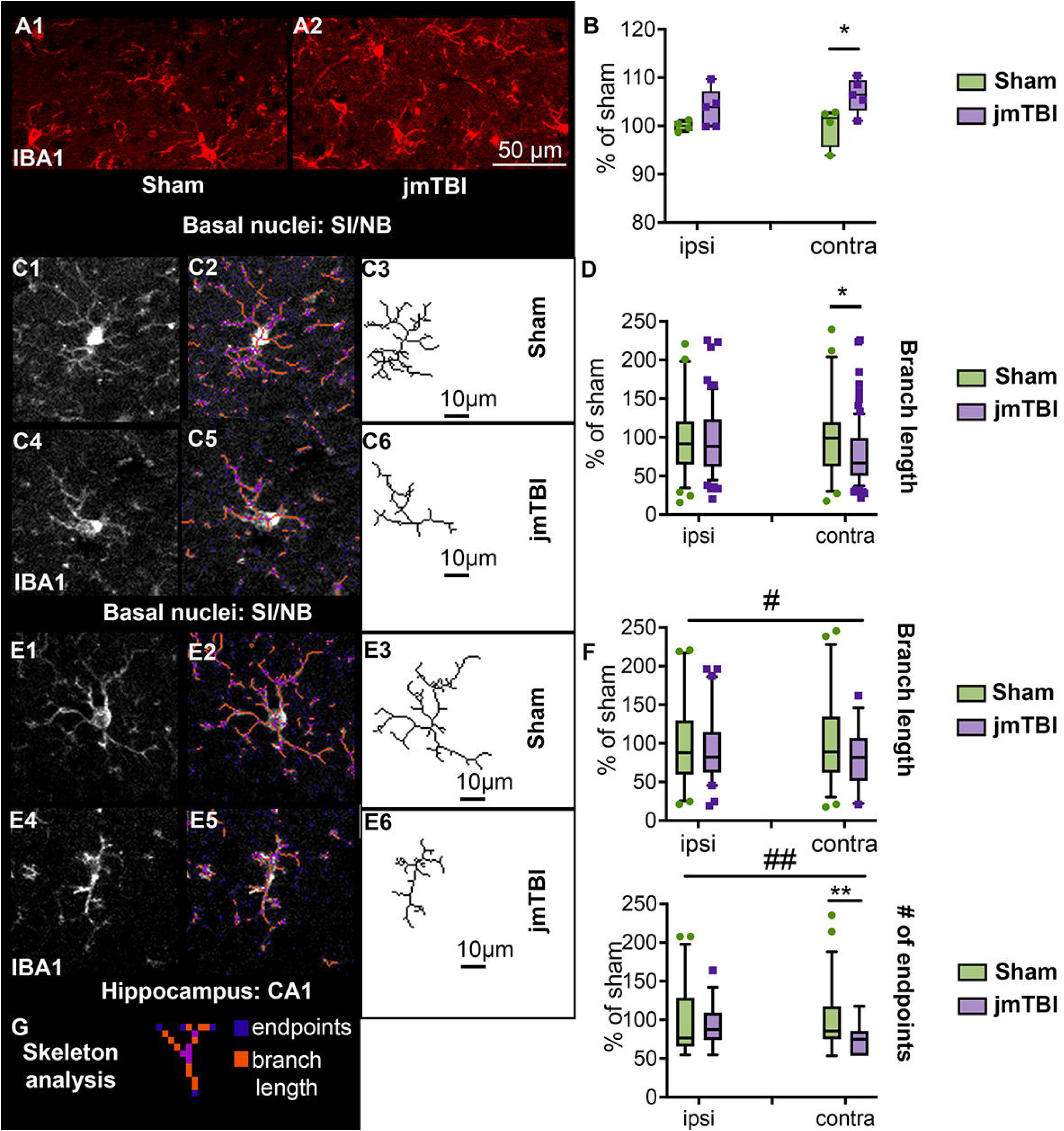
Microglial morphology was altered in SI/NB and hippocampus 12m after jmTBI. A. Increased microglial density in SI/NB after jmTBI (A2) compared to shams (A1). B. Iba-1 staining (mean OD, as %sham) in the contralateral SI/NB were significantly increased in the jmTBI compared to the sham cohorts, . Two-way ANOVA with Sidack post-hoc test. Data expressed as mean+SEM. * P>0.05. C. Individual microglia from sham (C1-3) and jmTBI (C4-6) and their respective tagged skeletons and isolated skeleton structures from the SI/NB D. Quantification of microglial skeleton parameters revealed decreased branch length in the contralateral SI/NB of jmTBI mice. E. Microglia from the CA1 and their respective tagged skeletons and isolated skeleton structures in the in sham (E1-3) and jmTBI (E4-6) mice. F. Microglia from jmTBI mice have decreased branch length and endpoints in CA1, especially in the contralateral side. Two-way ANOVA (# indicating jmTBI vs sham overall in ipsi and contra) with Sidack post-hoc test (* indicating jmTBI vs sham differences within the ipsi or contra side). Data expressed as mean+SEM. * or #P<0.05, ** or ##P<0.01 G. Illustration of skeleton analysis parameters. (passed Shapiro-Wilk normality test)

### Long-term cognitive deficits 12months after TBI

jmTBI and sham groups were tested in the elevated plus maze (Figure 8A), open field (Figure 8B) and sucrose preference (Figure 8C) tests at 6 and 12 months post-injury to assess behavioral alterations.

**Figure 8:**
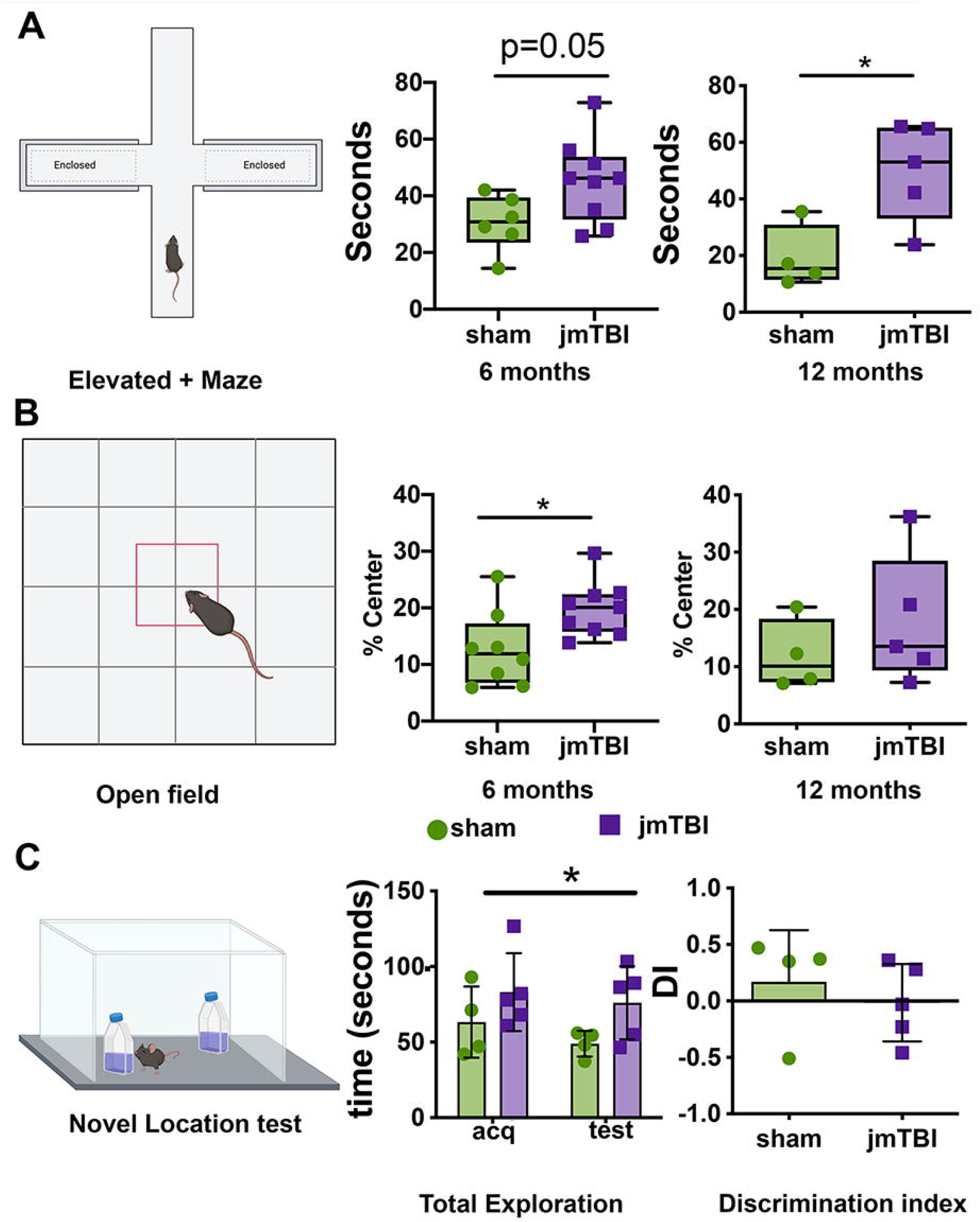
Impaired behavioral outcomes are found at 6 and 12m after jmTBI. A. Illustration of the elevated plus maze test with enclosed and open arms. jmTBI mice showed increased time in the open arms at 6 (p=0.05) and at 12 months (p<0.05) compared to sham mice, suggesting decreased anxiety like behavior. B. Illustration of the open field test performed at 6 and 12 months. Similar to the elevated plus maze test, the jmTBI group spent significantly more time in the center of the maze compared to shams at 6 (p<0.05) and 12 months (p=0.38). C. Illustration of the novel object location test, in which jmTBI mice displayed significantly higher exploration of both new and old locations. (Two-way ANOVA with Sidack post-hoc test, # indicates sham vs TBI differences). There were no significant differences between groups in the preference for a new location (discrimination index, DI).

In the elevated plus maze, jmTBI mice spent more time in the open arms compared to shams at 6 months p=0.05; t=2.152, df=13) and 12 months post-injury (p=0.0192; t=3.028, df=7) (Figure 8A). In the open field test, jmTBI mice spent more time in the center at 6 months but not at 12 months post-injury (Figure 8B). Finally sucrose preference testing showed a significant increase of consumption of the sweet solution drinking in jmTBI group at 12 months (supplementary figure 4).

The jmTBI mice exhibited decreased exploration of new locations in the novel object location test (Figure 8C), consistent with altered learning and memory. In addition, Morris Water Maze (MWM) testing was performed at 12 months post-injury (Figure 9). Both sham and jmTBI groups performed identically during the cued test, being able to find the visible platform with similar swimming speeds (data not shown), confirming no major alteration in visual and motor functions at 12 months after injury. However, jmTBI mice exhibited spatial learning deficits, with significantly increased cumulative distance to find the hidden platform compared to shams (Figure 9A). At 24h after the last trial, the probe test was performed to assess the spatial memory (Figure 9B) where jmTBI mice did not show any quadrant preference, while sham animals significantly preferred the platform quadrant (Figure 9B). Together, these results would suggest memory deficits in jmTBI mice at 12 months post-injury.

**Figure 9:**
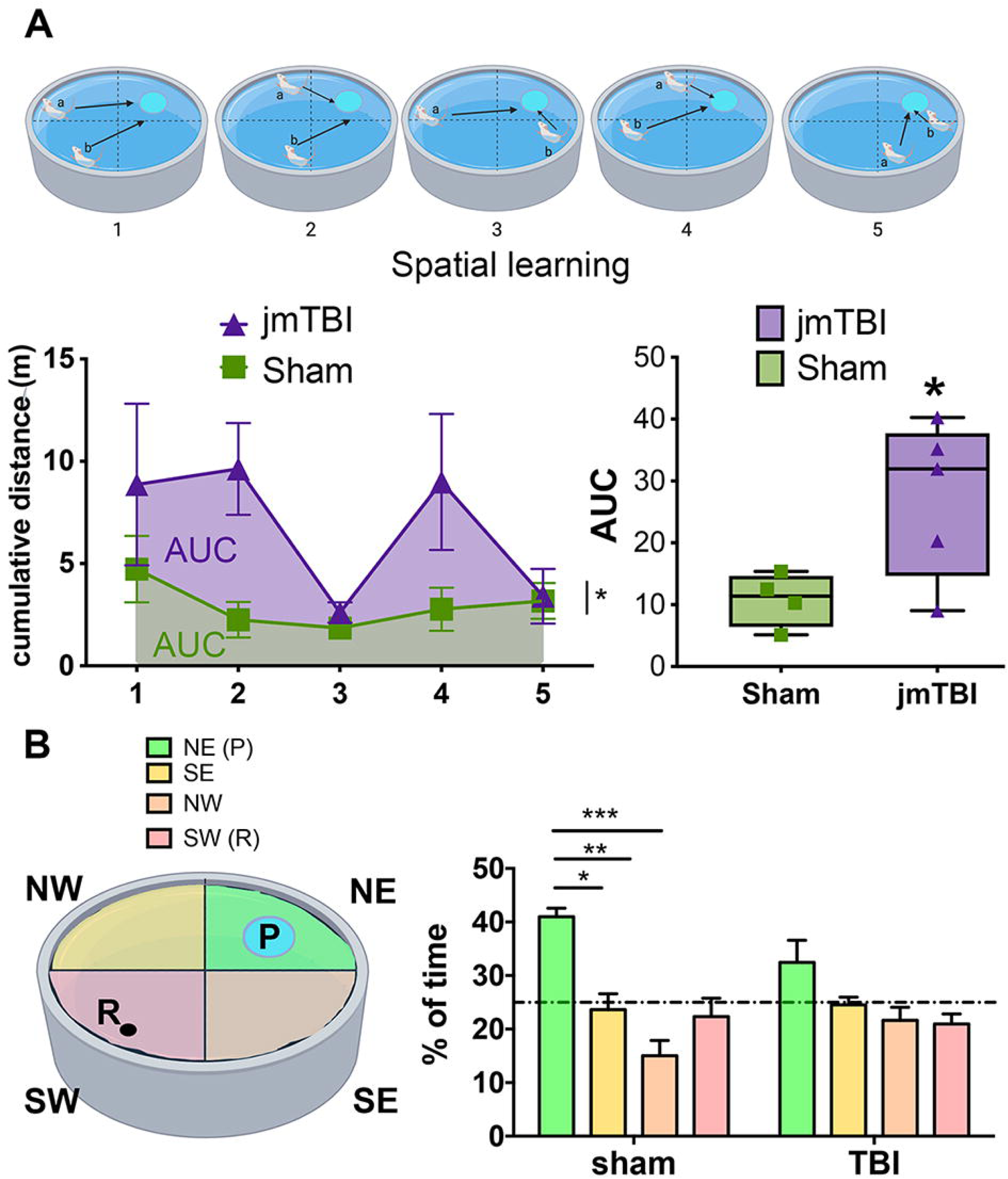
Impaired learning was found 12m after jmTBI. A. Illustration of the platform positions (blue) and two release positions for the animals (a, b arrows) within the Morris water maze (MWM, top panel). Spatial learning during the MWM was impaired in the jmTBI cohort, with increased cumulative distance to the platform over the trials during the learning task. B. In the probe test for spatial memory, the jmTBI mice had no significant preference for the platform (P) quadrant (in green) (left panel), whilst sham animals spent significantly more time in the platform quadrant compared to the other quadrants (right panel). The release point of the mice is indicated with R. Two-way ANOVA with Sidak post-hoc test. Data expressed as mean+SEM. *P<0.05, **P<0.01, ***P<0.001

### Relationships between DTI modifications and learning/memory deficits

We next evaluated how the altered DTI metrics observed in both SI/NB and hippocampus from mice after a single jmTBI were correlated to the performance of these animals in tests involving spatial learning, memory and discrimination.

We found that FA values in the ipsilateral CA1 were significantly correlated to the cumulative distance during the spatial learning task (Figure 10A; r^2^=0.6313, p=0.0105).

**Figure 10:**
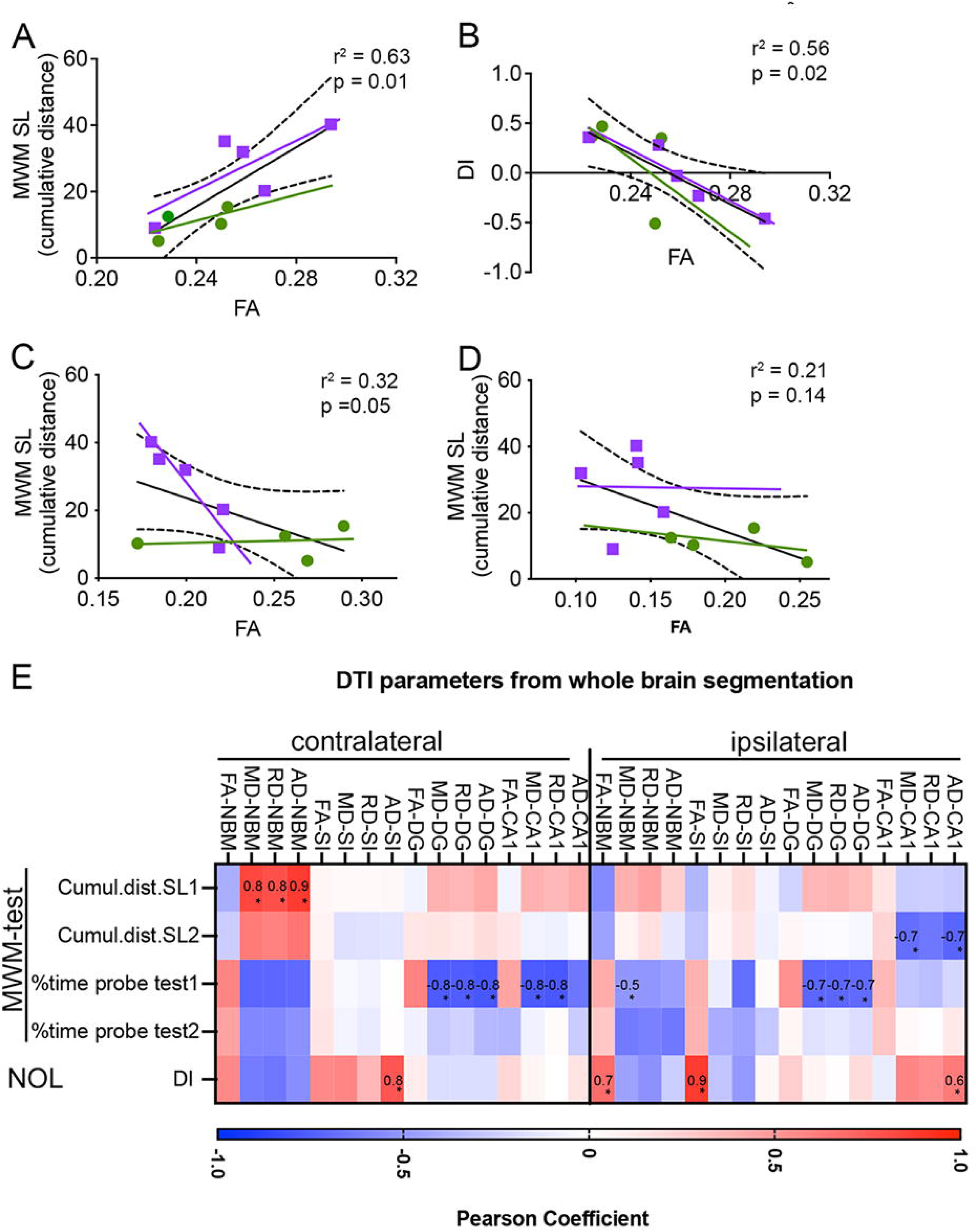
Correlations between DTI derived fractional anisotropy from manual and whole brain segmented analyses and behavior. Green circles represent individuals from the sham group and purple squares are the individuals from the jmTBI group. A. Higher FA in the ipsilateral CA1 region correlated with poor learning (longer cumulative distance) during spatial learning task 1 (SL, A) of the MWM. B) Increased FA in the ipsilateral CA1 region negatively correlated with worse memory (lower DI) during the NL test. C, D. Decreased FA in the ipsilateral (C) and contralateral (D) SI/NB postively correlated with poorer memory (lower DI) during the NL test. Linear regression. Dashed lines represent intervals of confidence. E) Pearson correlation coefficients between spatial memory test outcomes and DTI values from whole brain segmentation analyses for ipsi and contralateral CA1, DG, SI and NBM. The heatmap shows in red positive and in blue negative correlations, with significant differences inidcated by an asterisk (p<0.05). Several DTI values in the 4 ROIs ipsi and contra are correlated with behavioral scores.

We also noted that FA values in the ipsilateral CA1 were inversely correlated to the discrimination index of the novel object location (Figure 10B; r^2^=0.5636, p=0.0197; see Supplementary Table 2). In contrast, FA values in both ipsi and contralateral SI/NB were positively correlated to the discrimination index of the novel object location (ipsilateral: Figure 10C, r^2^=0.5057, p=0.0315; contralateral: Figure 10D, r^2^=0.5886, p=0.0158).

These results suggest that FA values within the vulnerable regions could represent a predictive metric for the outcome on spatial learning and discrimination capacities, particularly later in life.

## Discussion

While the consequences of a mild traumatic brain injury sustained during adulthood have been described, the behavioral and neurophysiological consequences of such trauma sustained early in development are still debated. The results of the present study suggest that in an animal model of juvenile concussion the emergence of cognitive decrements months after the trauma correlated with hippocampal and SI/NB MRI microstructural alterations. Furthermore, these alterations paralleled histological analysis of neuronal and glial cells which demonstrated significant molecular and morphological alterations to the neurovascular unit. Notably such neurovascular unit modifications share similarities with described in other neurodegenerative disorders [44].

### Diffusion neuroimaging of brain structure correlates with behavioral decrements

Diffusion alterations in white matter structures have been related to cognitive decline following pediatric TBI [37, 56]. However, microstructural alterations in grey matter have not been well studied in this context. In our model, the cognitive impairments at 12 months post-injury significantly correlated with the altered diffusion properties of the hippocampus and the basal forebrain, both of which are known to play a critical role in memory. This would suggest that DTI derived metrics may represent a non-invasive biomarker to identify brain tissue properties post-mTBI.

The distinct patterns of change in DTI metrics between brain regions (hippocampus vs. SI/NB) suggest apparent differences in microstructural and molecular profiles. In conjunction with our immunohistochemistry findings, the following inferences can be made. Firstly, decreased FA and AD, with increased NF200 expression in the basal forebrain would be consistent with axonal injury where the observed beading pattern of staining reflects injured fibers [10, 46]. Alternatively or in conjunction, AQP4 decreases could also contribute to reduced water diffusion as observed in decreased FA and AD. Secondly, the hippocampus exhibits an opposite phenotype to the SI/NB, where an FA increase parallels increases in AQP4 expression. Neuronal decrements are reflected by a decrease in both number (NeuN staining) and structure (APP staining). Previously in a CCI model, increases in GFAP staining were correlated to increased FA [9] and the expression level of AQP4 has been previously shown to modify the DWI signal [6]. Thus, based on our findings we speculate that in the hippocampus increased GFAP and AQP4, accompanied by loss of neurons, may contribute to increase of grey matter anisotropy [9].

More detailed whole-brain analysis of hippocampal subregions and basal forebrain circuit components revealed similar insights into the long-term microstructural alterations at 12 months following jmTBI. The significant findings from manual region of interest analysis in DTI metrics (Figure 2) and the lack of significant results from whole brain atlas derivations (Table 1) initially appear at odds. However, the manual DTI measures were extracted solely from the dorsal hippocampal regions (as illustrated in Figures 1, 2) whereas the atlas-based approach encompassed the entire hippocampus, dorsal to ventral regions. The curvilinear nature of the hippocampus results in different diffusion metrics dependent upon where in the hippocampus measures are made, thus averaging these variations resulted in no significant findings from whole hippocampal DTI measures. However, we found strong correlations between whole hippocampi DTI values and behavioral measures, consistent with our manual dorsal hippocampus analyses (Figure 10).

### Region-dependent neuronal alterations

The chronic consequences of jmTBI are different between basal forebrain and hippocampus regions using MRI and histological approaches. Alterations in the hippocampus and the basal forebrain have been previously described in clinical [32] and preclinical [19, 26] moderate/severe pediatric or juvenile TBI. The increased NF200 expression in the basal forebrain suggests alterations in neuronal fibers [46]. Similarly APP staining showed differences in neuronal processes injury in the hippocampus suggesting possible swelling and beading. Our data are in accordance with previous reports in moderate/severe TBI that report an early (1 to 3 weeks) loss of cholinergic neurons within the basal forebrain, a cholinergic structure projecting to the subcortical structures and important in cognitive functions after brain injury [1, 18, 24, 48]. Further, our preclinical data are in line with long term neurodegeneration observed in many patients with CTE or repetitive mTBI patients [40, 41]. Basal forebrain cholinergic neurons project to hippocampal structures [7] and alterations of basal forebrain neuronal fibers likely contribute to the loss of hippocampal neurons reflected by decreased NeuN staining via a neurodegenerative mechanism [47]. Finally, neuronal alterations (loss of cell bodies and fiber alterations) in CA1 and DG has been associated with local neuroinflammation leading to increased GFAP and AQP4 staining, microglia activation, and vascular alterations. There is a striking similarity of our findings to those found in TBI related dementia [47].

### Region-dependent neurovascular alterations

Early after jmTBI, neurovascular changes have been observed in various brain regions including corpus callosum [57], cortex [15, 27] and hippocampus [15]. These early alterations can be transient similar to cortical hypoxia (first 24h), accompanied by altered vascular reactivity and BBB hyperpermeability [27, 57]. These acute altered vascular morphological changes and increased GFAP expression have been observed up to 1 month after the jmTBI [15, 27]. Our current findings confirm that neurovascular phenotypic transformations are still present at 12 months after mild jTBI but these differences are brain region dependent. First, the opposite findings of decreased GFAP- and AQP4-staining levels on astrocytes in the basal forebrain contrasts with no change for GFAP- and increases for AQP4 staining intensities observed in the hippocampus. A redistribution of AQP4 away from astrocytic endfeet has been reported after severe adult TBI and in animals developing post-TBI epilepsy [66] which contrasts to the increased level of expression of AQP4 on astrocyte endfeet in contact with blood vessels in the hippocampus in our jmTBI model.

Region-dependent changes in AQP4 (increase vs. decrease) have been previously observed after stroke in white vs. grey matter [64] and sub-cellular AQP4 alterations may be linked to accelerated aging and cognitive impairments through its role in perivascular brain clearance of protein aggregates [77]. Also AQP4 levels are reported toincrease with age [23, 77] and these increases are associated with tau [34] and β-amyloid [61] deposition. AQP4 levels are increased in Alzheimeŕs disease in human samples [77]. While the linkage of high levels of AQP4 to Alzheimeŕs disease are tantalizing, it may not be a primary mechanism of pathology as AQP4 deletion has also been associated with increased protein accumulation, most notably after TBI [28]. A caveat is that the total amount of AQP4 is not as relevant for pathology progression but rather where AQP4 is localized. In support of this hypothesis, decreased perivascular AQP4 correlated with increased β-amyloid accumulation in humans [77] and in mice [35, 75]. Therefore, decreased perivascular AQP4, as observed in neurodegenerative aging models, could be closer to the pathology we observed in the basal forebrain compared to that of the hippocampus after jmTBI. In fact, reallocation of AQP4 away from the perivascular endfeet has also been observed in post-stroke dementia [12], suggesting that it may be a common mechanism underlying different forms of cognitive dysfunction.

We found that blood vessels exhibited reduced vessel diameters in basal forebrain but not in hippocampus, with a higher number of blood vessels with a diameter under 5μm in agreement with our previous work [27]. Vascular modifications have been previously reported at 2 and 6 months post-jTBI that parallel β-amyloid accumulation and cognitive decrements [29, 49]. This would suggest that vascular alterations may be a common pathological landmark of a neurodegenerative processes; where decreased blood vessel diameters has been described in neurodegenerative disease models [44] and Alzheimer’s disease [67], leading to modified cerebral perfusion.

Our observation that at 12months after jmTBI there was increased microglia activation (increased IBA1 with more protracted morphology) in both hippocampus and SI/NB that occured with similar blood vessel, astrocyte, and neuronal changes. In both the basal forebrain and the hippocampus, microglia acquired a more contracted and less ramified shape similar to the description made for other pre-clinical TBI models at early time points post injury [38]. In fact, microglial activation had been previously observed in the basal ganglia and the nucleus accumbens, during the first month after jmTBI, mediating addictive behavior [31]. In a model of adult mTBI, microglia morphological alterations in were also observed in cortical regions remote from the impact site [38]. These reports further support our findings that regions remote from the impact site are perturbed even after an early in life mTBI. While microglial activation was observed in both brain structures, hippocampal microglia exhibited more dramatic morphological alterations where the size of the processes and number of branches and endpoints were all significantly reduced within CA1, whereas in SI/NB only a decrease in processes size was observed. Activated microglia can have both beneficial and noxious effects, therefore a more detailed analysis of microglial markers would be required to identify the phenotypic expression of microglial functions in the context jmTBI.

The glial and vascular changes are comparable to the neuronal alterations in both structures with a change in the neuronal processes observed with NF200 and APP stainings, and a loss of neuronal cell bodies with NeuN staining in the hippocampus. MRI-derived brain tissue changes further support with the histopathologial alterations described in hippocampus and SI/NB remote from the site of the impact and at longterm post-injury.

### A single mild juvenile TBI event triggers longterm alterations

Our initial hypothesis that a single mTBI event during childhood could trigger long-term changes into middle age was confirmed in our jmTBI model. We found brain microstructural alterations correlated with altered behavioral outcomes (elevated plus maze, open field test, spatial learning and memory) and paralleled neurovascular changes in both the basal forebrain and hippocampus. From 2 to 12 months, various mouse behavioral repertoires were altered after jmTBI compared to sham group. Interestingly, injured animals spent more time in the center of the open field and open branches of the elevated plus maze associated with higher exploratory behavior in the novel object location. Altogether these results suggest that jmTBI animals are less anxious in developing novelty seeking behavior similar to a BLAST injury model [45]. This behavior is observed over time. We did not observe any differences in the cued performance on MWM (data not shown), strongly suggesting the specificity of the hippocampal-cholinergic alterations in spatial learning and memory. The spatial learning alterations at 12 months after jmTBI are also in accordance with previous work in mild adult closed head injury model of mTBI that showed decreased performance in learning at 12 months after injury [39] and similar to decrements observed in severe juvenile TBI models [30]. The age of the impact is important feature in rodent adolescent TBI models as spatial learning and memory dysfunctions are not always observed [13, 58]. Previous work showed that memory retention was affected after a single mTBI in adolescents but not in adults early after closed head injury [22]. In agreement with our findings, a single mTBI to the developing brain is sufficient to alter memory retention 12 months post-injury supporting the notion that the developing brain is more sensitive to injury and results in longterm alterations.

An emerging decrease in anxiety-like behavior is observed in jmTBI group with increased time spent in the center of the arena and in the open arms of the elevated plus maze test. Even then the direct comparison of our data to clinical observations is not straightforward. Neurophysiological changes in relation with post-concussive symptoms (PPCS) after pediatric mTBI have not been well characterized. Children with PPCS exhibited decreased cortical activation on fMRI in working memory tasks at one-month post-injury [33]. Similarly, resting state fMRI showed hyperconnectivity and hypoconnectivity in sport related concussions that were associated with altered memory performance up to 21days [43]. These clinical data link MRI changes with cognitive outcomes. Similarly, we observed a correlation between neuroimaging changes with behavioral outcomes and histology alterations in our jmTBI model up to 12 months after the injury. Some clinical studies suggest a recovery at longterm in children who endured a concussion 2.5 years prior to study [8]. This difference brings the question of how we can compare the elapsed time after injury in rodent model and human brain pathology. Longterm alterations could be in future treated, as suggest by a recent clinical work proposing to use transcranial direct current stimulation to treat PPCS in children [52].

Clearly differences between adult brains and developing brains supports the hypothesis that the outcome after pediatric TBI leads to more severe outcomes, particular later in life, compared to adults for the same degree of TBI severity [21]. In line with our previous and current preclinical findings, clinical studies are now reporting cognitive deficits 10 years after an early in life brain injury [3, 4].

### Study Limitations

There are several limitations in our study. A limitation is the small number of animals at the end of the 12 months post-injury injury that was a result of extracting mice at selected time points for ex vivo MRI and correlative histological analyses. Moreover, given known sex differences this study should be replicated examining how sex modulates mTBI outcomes. To address these gaps, we have an ongoing study with a larger cohort comprised of male and female mice with various timepoints to monitor the temporal evolution using imaging, blood biomarkers and histology. The histological findings reported here will be confirmed in future research using RNA and Western blot analyses. Similarly, we cannot exclude that other brain regions exhibit DTI changes, however we decided to focus on the cholinergic system known to be involved in the memory system.

## Conclusions

We report for the first time the emergence of long-term cognitive deficits that correlate with tissue microstructural (DTI) modifications and are coincident by robust neurovascular pathophysiology (figure 11). Not surprisingly, the pathology was regionally dependent and are consistent with an ongoing slow but progressive degenerative process(es) contributing to a potential loss of projections from the basal forebrain toward the hippocampus. Most significantly, a single mTBI during vital brain developmental periods are sufficient to evoke long-term functional consequences that are seen in middle age, affecting cognitive ability and potentially impacting cognitive reserve.

**Figure 11:**
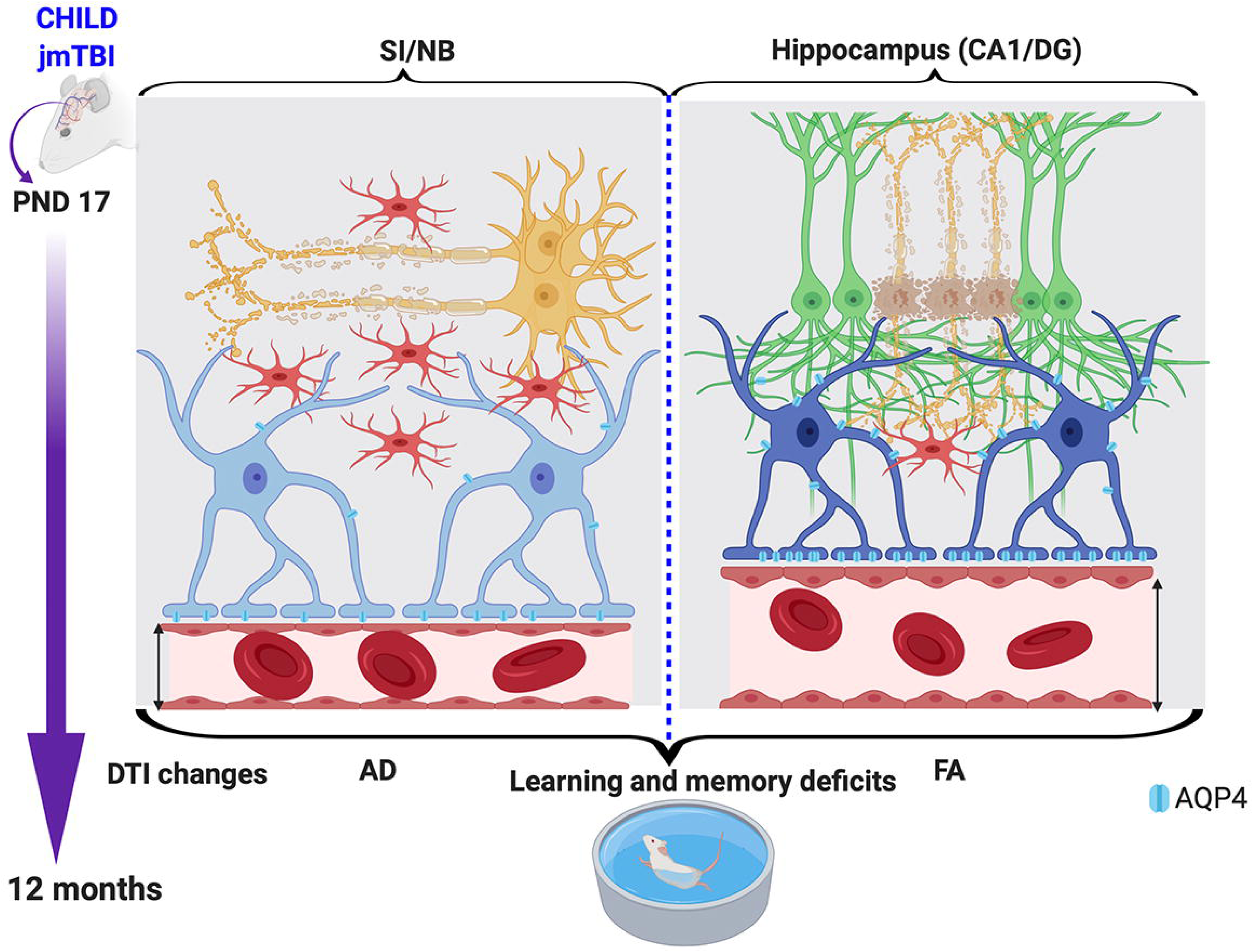
Summary of the consequences following a single jmTBI that leads to accelerated aging (12 months post-injury). In the SI/NB, there was reduced astrogliosis (GFAP staining) with reduced levels of AQP4 in astrocyte processes and at perivascular endfeet. This was accompanied by increased microgliosis (levels of Iba1 staining and morphological alterations), decreased vessel diameter and altered axonal staining (NF200). In the hippocampus, AQP4 levels were increased, concomitant with increased microglial activation (morphological changes in Iba1-stained cells). Neurodegeneration (decreased NeuN) was observed in the ipsilateral hippocampus. All these changes were accompanied by DTI alterations that correlated with memory and learning deficits.

## Declarations

The authors have no financial conflicts to report.

## Ethics approval

All animal procedures were carried out in accordance with the European Council directives (86/609/EEC) and the ARRIVE guidelines.

## Consent for publication

n.a.

## Availability of data and material

The datasets used and/or analyzed during the current study available from the corresponding author on reasonable request.

## Competing interest

Authors declare no competing interests.

## Funding

This research project was funded by Eranet Neuron TRAINS, Neuvasc, ANR Nanospace and RO1, NINDS.

## Authors’ contributions

BRG performed TBIs, analysis of behavioral data, MRI data acquisition and manual ROI analysis, histology and image analysis, prepared the manuscript and contributed to experimental design. MLF performed behavioral tests and contributed to the analysis of behavioral data. JL performed the whole-brain analysis and contributed to manuscript writing. CD, MC, and SCG critically revised the manuscript, JB and AO contributed to experimental design, data analysis, manuscript writing and revision. JB is leading author.

## Figure legends

**Supplementary figure 1.**
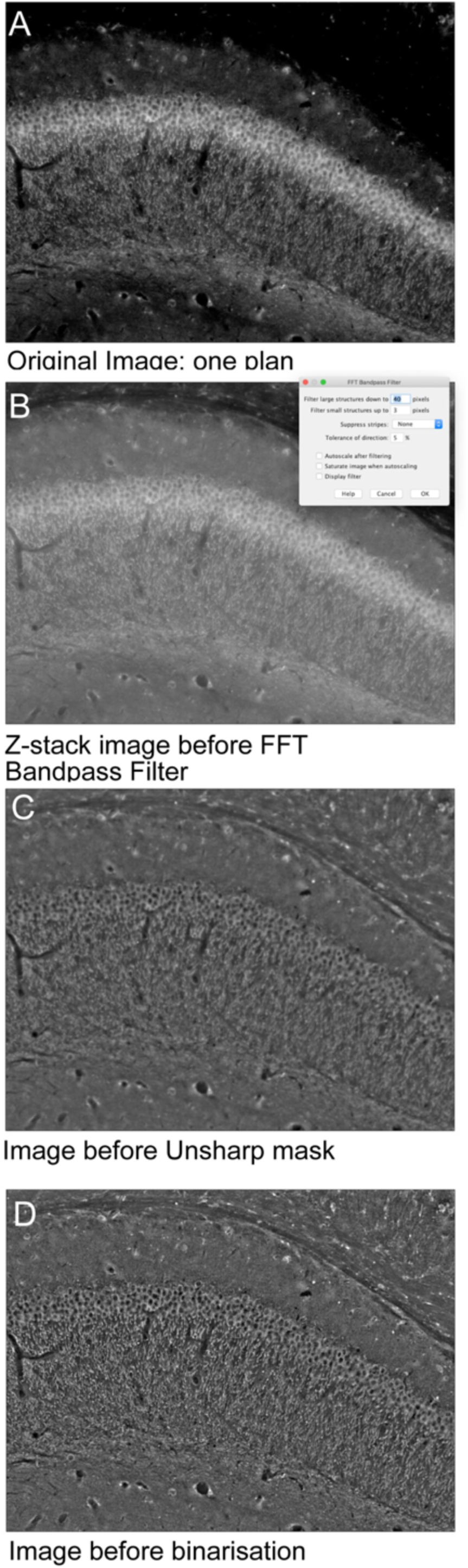
Image analysis process. Using Z-stack function in FIJI software, stack images (A) were reconstructed in one plane (B). Then, reconstructed image was treated with “Bandpass” filter without autoscale after filtering (C). The result image was treated with “Unsharp mask” (D) and then binarised.

**Supplementary figure 2.**
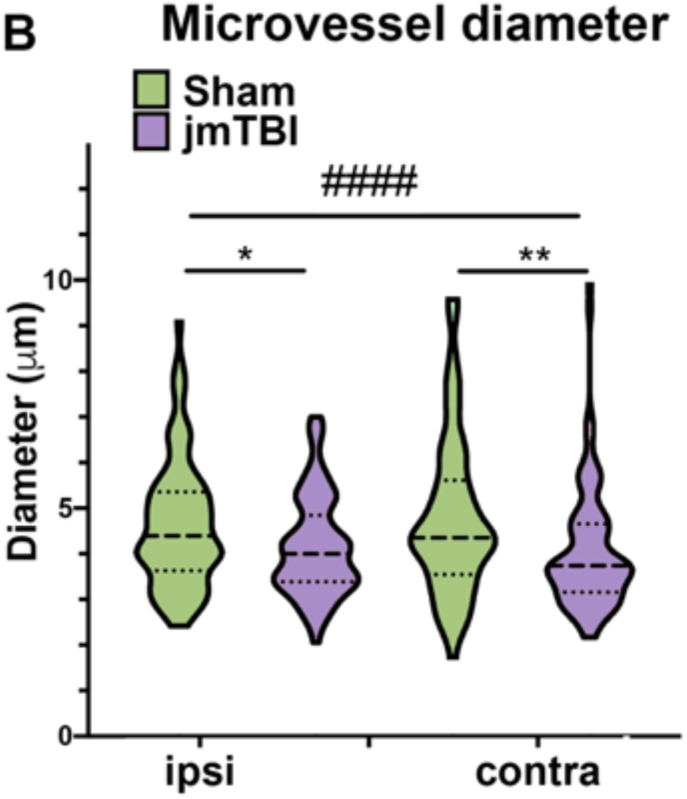
Decreased vascular diameter is observed in the SI/NB 12m after jmTBI. The reduction in vessel diameter in jmTBI compared to sham mice was significant in both ipsilateral and contralateral sides of the SI/NB. Two-way ANOVA (# indicating global jmTBI vs sham difference) with Sidack post-hoc test (* indicating jmTBI vs sham differences in the ipsilateral or contralateral side). Data expressed as mean+SEM. **P<0.01, ####P<0.0001

**Supplementary figure 3.**
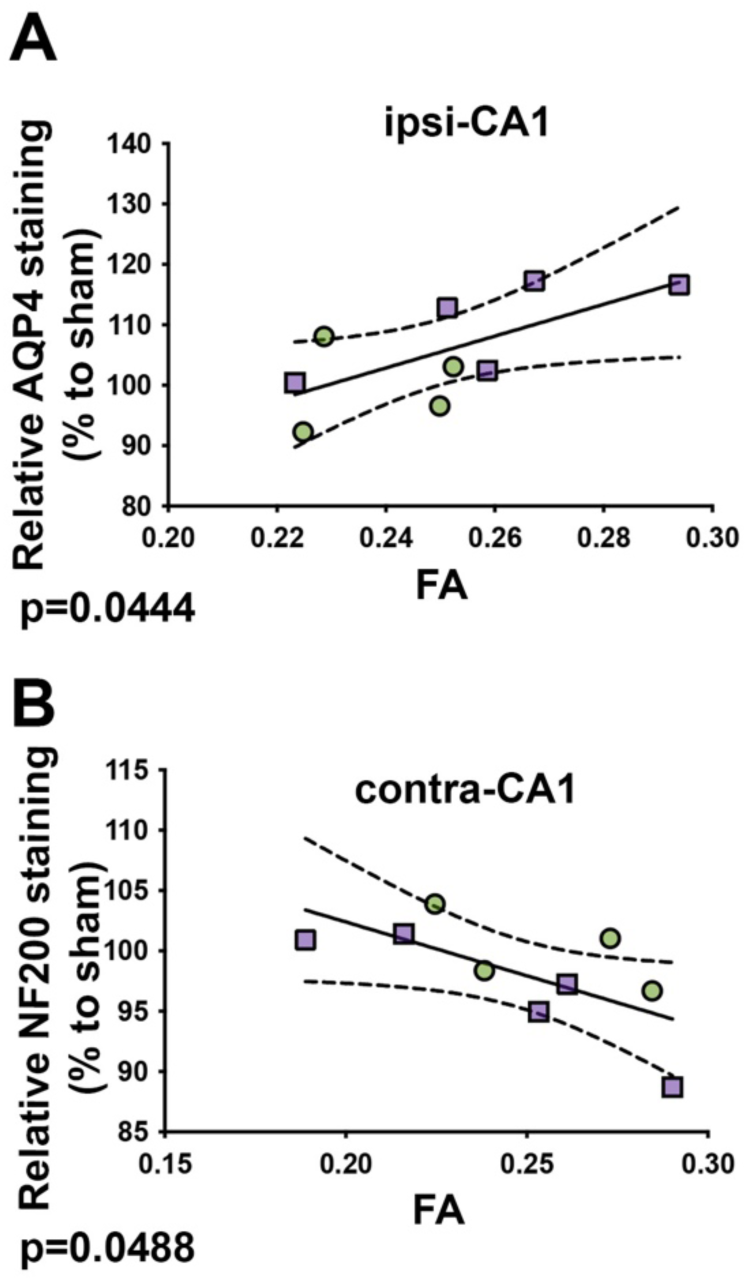
Correlation between immunolabeling and manual analysis of regional DTI changes

**Supplementary figure 4.**
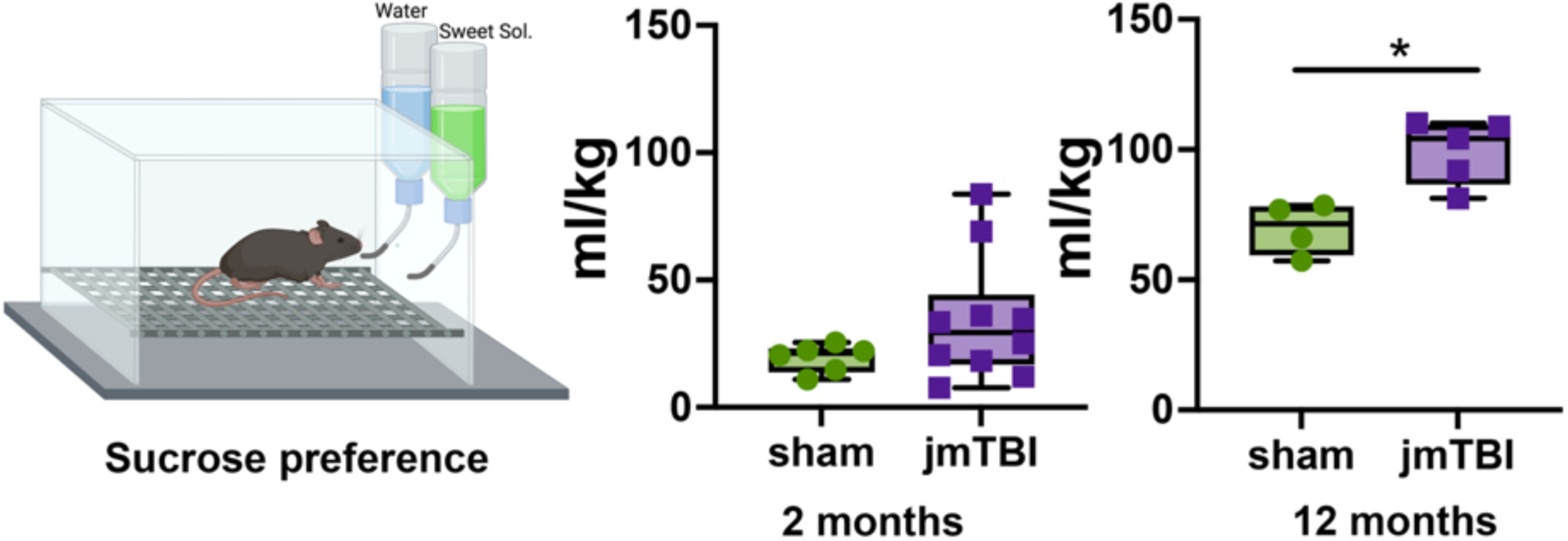
The sucrose preference test was performed at 2 and 12 months. A significant increase in sucose consumption was observed at 12 months compared to the sham (p<0.05).Two-way ANOVA with Sidak post-hoc test. Data expressed as mean+SEM. *P<0.05

**Supplementary table 1.** Imaging parameters.

|  | *In vivo* MRI |  |
| --- | --- | --- |
| Seq. Type | T2 (Bruker: MSME) | DTI (Bruker: DtiEpi) |
| TR (ms) | 3290 | 1000 |
| TE (ms) | 7 | 30 |
| NEX | 2 | 1 |
| FOV (mm) | 16x12.8 | 16x12.8 |
| Matrix | 162X128 | 162x128 |
| Slices (n) | 24 | 48 |
| Slice Thickness (mm) | 0.533 | 0.266 |
| Slice Interval (mm) | 0 | 0 |
| Echos | 25 | 1 |
| Flip Angle | 0 | 90 |
| Resolution (mm) | 0.098 x 0.1 | 0.098 x 0.1 |
| Number of directions | n.a. | 21 |

**Supplementary table 2.** DTI values and group comparisons from manual DTI analysis. Significant differences are higlighted in red and in bold. FA_fractional anisotropy, AD_axial diffussivity, RD_radial diffussivity, MD_mean diffussivity. *p-values correspond to Sidak post-hoc test following Two-way ANOVA, sham vs jmTBI.

| DTI | ROI |  | stats | sham | jmTBI | DTI | ROI |  | stats | sham | jmTBI |
| --- | --- | --- | --- | --- | --- | --- | --- | --- | --- | --- | --- |
| FA | DG | ipsi | mean | 0,2800 | 0,2466 | RD | DG | ipsi | mean | 0,3487 | 0,3785 |
|  |  |  | p* |  | **0,0477** |  |  |  | p* |  | 0,4524 |
|  |  | contra | mean | 0,2754 | 0,2705 |  |  | contra | mean | 0,3582 | 0,3761 |
|  |  |  | p* |  | 0,9635 |  |  |  | p* |  | 0,8014 |
|  | CA1 | ipsi | mean | 0,2389 | 0,2589 |  | CA1 | ipsi | mean | 0,4569 | 0,4138 |
|  |  |  | p* |  | 0,2818 |  |  |  | p* |  | 0,2104 |
|  |  | contra | mean | 0,2552 | 0,2419 |  |  | contra | mean | 0,4295 | 0,4544 |
|  |  |  | p* |  | 0,7665 |  |  |  | p* |  | 0,6567 |
|  | SI/NB | ipsi | mean | 0,2468 | 0,2008 |  | SI/NB | ipsi | mean | 0,2889 | 0,2639 |
|  |  |  | p* |  | 0,2526 |  |  |  | p* |  | 0,5322 |
|  |  | contra | mean | 0,2575 | 0,2424 |  |  | contra | mean | 0,2603 | 0,2580 |
|  |  |  | p* |  | 0,8477 |  |  |  | p* |  | 0,9943 |
| AD | DG | ipsi | mean | 0,5459 | 0,5544 | MD | DG | ipsi | mean | 0,4145 | 0,4371 |
|  |  |  | p* |  | 0,9377 |  |  |  | p* |  | 0,6241 |
|  |  | contra | mean | 0,5572 | 0,5735 |  |  | contra | mean | 0,4245 | 0,4419 |
|  |  |  | p* |  | 0,7459 |  |  |  | p* |  | 0,7797 |
|  | CA1 | ipsi | mean | 0,6328 | 0,5976 |  | CA1 | ipsi | mean | 0,5155 | 0,4751 |
|  |  |  | p* |  | 0,3589 |  |  |  | p* |  | 0,2486 |
|  |  | contra | mean | 0,6125 | 0,6290 |  |  | contra | mean | 0,4905 | 0,5126 |
|  |  |  | p* |  | 0,7424 |  |  |  | p* |  | 0,6724 |
|  | SI/NB | ipsi | mean | 0,4236 | 0,3604 |  | SI/NB | ipsi | mean | 0,3338 | 0,2961 |
|  |  |  | p* |  | **0,0294** |  |  |  | p* |  | 0,2219 |
|  |  | contra | mean | 0,3919 | 0,3798 |  |  | contra | mean | 0,3042 | 0,2986 |
|  |  |  | p* |  | 0,8440 |  |  |  | p* |  | 0,9638 |

## Notes

### Competing Interest Statement

The authors have declared no competing interest.

